# Effects of curcumin nanoformulations on cellular function in Niemann-Pick disease type C astrocytes

**DOI:** 10.1101/135830

**Authors:** Emily Maguire, Luke J. Haslett, Joanne L. Welton, Helen Waller-Evans, Jule Goike, Emily H. Clark, Harry R. Knifton, Ravin Shrestha, Kim Wager, Richard Webb, Emyr Lloyd-Evans

**Affiliations:** School of Biosciences, Sir Martin Evans Building, Cardiff University, Museum Avenue, Cardiff, CF10 3AX. United Kingdom; Department of Biomedical Sciences, Cardiff Metropolitan University, Llandaff Campus, Western Avenue, Cardiff, CF5 2YB, United Kingdom

**Keywords:** Niemann-Pick, curcumin, nanoformulation, lysosome, therapy

## Abstract

Niemann-Pick disease type C1 (NPC disease) is a neurodegenerative multi-lipid lysosomal storage disease caused by mutations in the *NPC1* gene presenting with reduced lysosomal Ca^2+^ signalling and inhibited late endosome-lysosome transport. Elevating cytosolic Ca^2+^ levels in NPC cells has been shown to reduce lysosomal lipid storage. Treating *Npc1^-/-^* mice with the Ca^2+^ modulator curcumin led to reduced lipid storage, improved life expectancy and function. These studies led to reported utilisation of curcumin supplements by NPC disease families despite there being no clinical evidence of benefit and a report indicating no benefit of nanoformulated curcumin in *Npc1^-/-^* mice. The aim of this study was to determine whether various commercially available curcumin nanoformulations were capable of reproducing the findings obtained with unformulated pharmaceutical grade curcumin. We compared seven curcumin nanoformulations in *Npc1^-/-^* mouse astrocytes. All the nanoformulations elevate cytosolic Ca^2+^ levels but only two lowered lysosomal lipid storage. Importantly, some caused elevations in NPC lysosomal storage and/or decreased cellular viability. Although this is an *in vitro* study, our findings suggest that care should be taken when contemplating the use of curcumin supplements for NPC disease.

## Background

NPC disease is a neurodegenerative lysosomal storage disease characterised by progressive ataxia, hallmarks of Alzheimer disease, dystonia and hepatosplenomegaly^1^. At the cellular level the disease is characterised by accumulation of multiple lipids including cholesterol, sphingosine, glycosphingolipids (GSLs), lyso-(bis)phosphatidic acid (LBPA) and sphingomyelin within late endosomes and lysosomes^1-3^. The disease is caused by mutations in either the *NPC1* or *NPC2* genes with the *NPC1* gene encoding a 13 transmembrane domain protein (NPC1) residing within the limiting membrane of the lysosome^1^. Whilst the exact function of this protein remains unknown it has been shown to bind to cholesterol and to have greatest homology to the resistance nodulation division (RND) family of bacterial multi-substrate efflux pumps^1,4^. At present, the only approved therapy in Europe for NPC disease is miglustat^5^, which reduces GSL biosynthesis and storage^6^, and if given early enough can slow disease progression but does not reverse the disease course^7^.

A major contributing factor to the accumulation of lipids in NPC1 disease cells is the retardation in endosomal transport including defective delivery of material to lysosomes^3^, defective fusion between late endosomes and lysosomes^2^, and defective transport of material between late endosomes/lysosomes and other organelles including the Golgi or the endoplasmic reticulum (ER)^1,8^. We have previously shown that an important contributing factor to these defects is the reduced nicotinic acid adenine dinucleotide (NAADP) mediated Ca^2+^ release from lysosomes^2^, a necessary trigger for mediating endocytic transport^9^. This endocytic trafficking defect in NPC cells can be overcome by inducing transient global elevations in cytosolic Ca^2+^ levels^2,10,11^. Using curcumin, a weak inhibitor of the sarco/endoplasmic reticulum (ER) Ca^2+^ ATPase (SERCA)^12^, significant Ca^2+^ release from the ER can be triggered leading to a transient large elevation in cytosolic Ca^2+^ that in turn corrected the endocytic trafficking defect and led to a reduction in lipid storage in NPC1 disease cells^2^. This benefit was caused directly by the elevation in cytosolic Ca^2+^ as it could be blocked using the Ca^2+^ specific chelator BAPTA-AM and had no effect on the reduced lysosomal Ca^2+^ levels^2^.

We observed similar benefit *in vivo* in the *Npc1^-/-^* mouse model with a reduction in lipid storage, an increase in life expectancy and improved function^2^. More recently, it has also been shown that curcumin treatment of the *Npc1^-/-^* mouse leads to ∼16% increase in lifespan (not dissimilar to the 25% increase observed with miglustat), improved performance on the balance beam^13^ and greater survival of Purkinje neurons with limited effect on inflammation, suggesting that the benefit is mediated by modulating Ca^2+^ levels rather than the antioxidant properties of curcumin^14^. Despite these studies it has been suggested that the concentration of curcumin that enters the blood and tissues is not high enough to modulate Ca^2^+^13^. A study in rats found that an oral dose of 2g/kg reached a maximum serum concentration of ∼4μM after 50 minutes^15^ and prolonged treatment of mice with 83mg/kg/day can maintain similar levels (∼1μM) within the brain^16^, concentrations that have been shown to have impact on behavior^17^. However, to boost the bioavailability of curcumin, and to reduce dosing, it is often encapsulated in a lipid mixture^13,18^. A study in mice demonstrated that 0.5% of a 5mg/kg injection of nanocurcumin was capable of entering the brain within 1h and achieving serum concentrations of ∼10μM^18^. Furthermore, a recent study has highlighted that different sized curcumin nanosupensions/nanoformulations occupy tissues at varying concentrations dependent on their particle size. For example, with initial plasma concentrations of around 5μM following an intravenous injection the highest concentration in the liver were observed with 200nm curcumin particles at 10 min post injection whereas in the brain the highest concentration was observed with 70nm particles at 20-30mins post injection^19^. These studies indicate that higher therapeutic concentrations of curcumin can be achieved using nanosuspensions and that tissue penetrance may be dependent on particle size. Interestingly however, no benefit on *Npc1^-/-^* mouse function was observed when curcumin was combined with a lipid carrier to increase bioavailability^13^. This suggests that either the lipid carrier interferes with the mode of action of curcumin or that particle size was sub-optimal.

It has come to our attention that some NPC disease patients are taking curcumin supplements^20^. Several of these curcumin supplements contain lipid mixtures consisting of phospholipids and fatty acids^15^. NPC disease cells are known to have defects in the efflux of phosphatidylcholine to apolipoprotein A1^21^, suggesting that some of these lipid mixtures could accumulate in NPC disease cells. Whilst some of the encapsulated curcumin would be delivered to cells of the intestine, which does not function normally in some NPC1 disease patients^22^, several nanoparticle vectors actually reduce clearance of the drugs they encapsulate resulting in accumulation in tissues such as the liver^23^, which is also affected in NPC1 disease^22^. As no information exists about the potential effects of these combined curcumin/lipid complexes on NPC1 disease cells, and as the previous study mentioned above did not observe any benefit when using curcumin complexed with a lipid vehicle^13^, we have undertaken a study to determine whether these curcumin nanoformulations are capable of reducing NPC lipid storage *in vitro* and whether their mode of action is in any way related to their nanoparticle size distribution.

## Methods

### Reagents

Unless otherwise stated, all reagents were obtained from Sigma-Aldrich, Dorset, UK.

### Cells

Primary astrocytes from post-natal day 1 *Npc1^+/+^* and *Npc1^-/-^* mouse were prepared as previously described^2^ and were maintained in culture as monolayers grown in Dulbecco’s modified Eagle’s medium (DMEM) supplemented with 10% foetal bovine serum (FBS) and 1% L-glutamine only at 37°C in a humidified incubator with 5% CO_2_. For all experiments, cells were seeded at equal densities of 5,000 or 20,000 cells on 8 well chamberslides (Ibidi, Thistle Scientific, UK) or Cellbind 96 well plates (Sigma-Aldrich, UK) respectively and left to adhere overnight prior to treatment with the curcumin formulations for the indicated times.

### Preparation and solubilisation of curcumin neutraceuticals

Curcumin supplements, summarized in Table 1, were purchased in multiple batches from commercial retailers. Three randomly selected capsules from each batch were opened, the powder weighed and solubilised in dimethylsulfoxide (DMSO, VWR, UK). We determined the curcumin content from the manufacturers’ stated ratios of curcumin to lipid carrier and bulking agents (Table 1, Fig. 1B) and generated a 10mM stock solution in each case. Each curcumin supplement was used at a final concentration of 30μM ensuring that the DMSO content in the treated cultures always remained at 0.3% v/v, appropriate DMSO and, where possible, lipid or bulking agent controls were also used.

**Table 1.** Properties of the curcumin nanoformulations.

| Referred to in this study as | Formulation | Producer | Reported lipid content | % Curcuminoid |
| --- | --- | --- | --- | --- |
| TRE | N/A (total root extract) | Solgar | No added lipids | 93% |
| CGM | Curcumagalactomannoside | Akay/Swanson | Fenugreek galactomannans | >35% |
| BCM95<sub>g</sub> | BCM-95 | Dolcas Biotech/Genceutic Naturals | Turmeric essential oil, lecithin, triglycerides, beeswax, sunflower oil | 95% |
| BCM95<sub>s</sub> | BCM-95 | Dolcas Biotech/LifeExtension | Turmeric essential oil | 95% |
| SLN<sub>A</sub> | MicroActive (Solid lipid curcumin particle) | Maypro Industries/Dr. Mercola | Medium chain triglycerides, polyglycerol oleate, sodium alginate | 25% |
| SLN<sub>L</sub> | Longvida (Solid lipid curcumin particle) | Verdure Sciences/AOR, Nutravene | Soy lecithin, palmitate, stearic acid | 23% |
| SLN<sub>M</sub> | Meriva (Solid lipid curcumin particle) | Indena/Dr.'s Best, Inc | Phospholipid | 37.2% |

**Figure 1.**
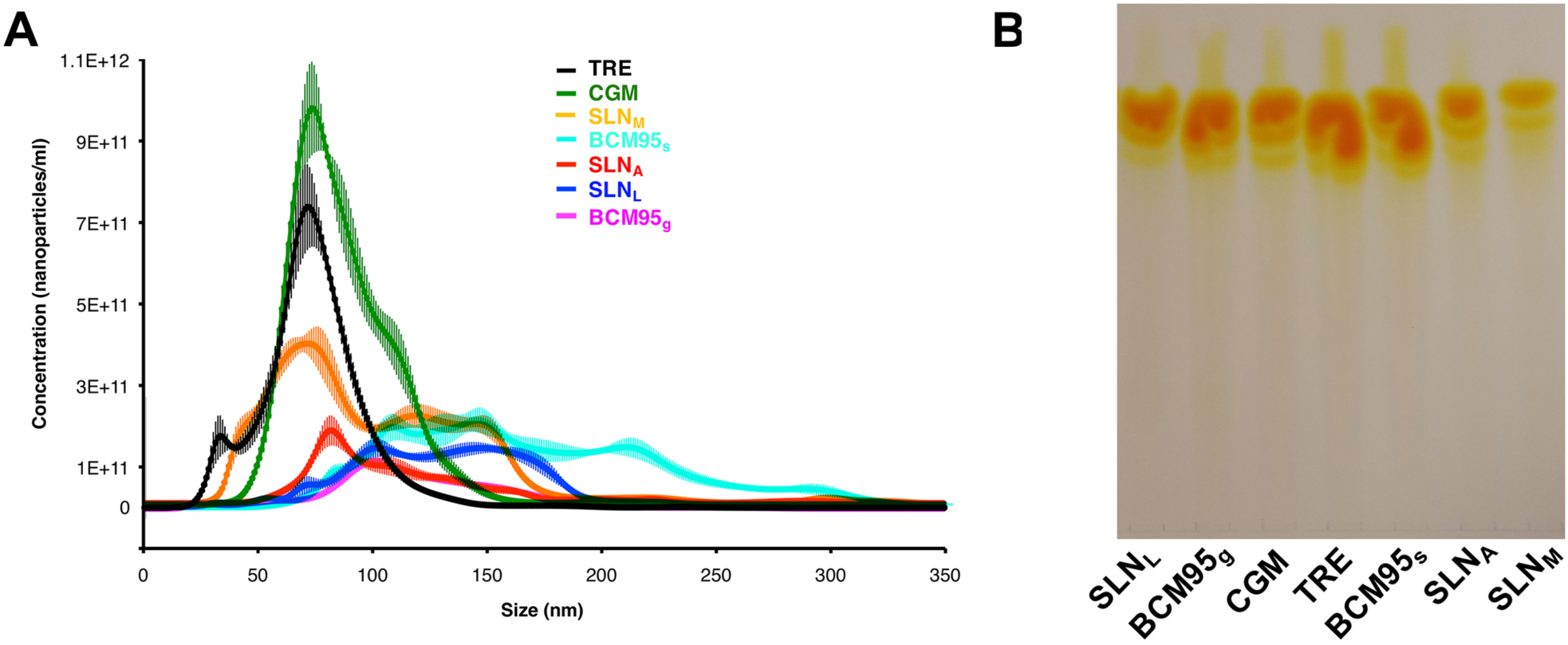
Curcumin nanoformulation particle size and curcumin content. (A) nanoparticle tracking was used to examine the size distribution of the nanoformulated curcumin. The mean ± standard deviation (s.d.) is shown for 5 separate measurements. (B) curcuminoid content (unstained) of each supplement separated by TLC. One whole capsule of formulated curcumin was solvent extracted and separated by TLC as described in materials and methods and separated by TLC, n=3.

### Cytosolic Ca^2+^ assays

Changes in cytosolic Ca^2+^ in response to curcumin formulations were measured using the membrane permeant acetoxymethyl (AM) derivative of the intracellular ratiometric Ca^2+^ indicator Fura-2,AM (Stratech Scientific, UK). Cells were loaded in DMEM as previously described^2^ with 5μM Fura-2AM and 0.025% Pluronic F127 for 1h at room temperature followed by 10min de-esterification of the AM ester group to release free cytosolic Fura 2. For all experiments, cells were bathed in an extracellular solution of Hank’s balanced salt solution (HBSS, Invitrogen, Paisley, UK) containing 1mM MgCl_2_, 1mM CaCl_2_ and 10mM HEPES (pH 7.2). Changes in cytosolic Ca^2+^ in response to the curcumin formulations were measured every second using 360nm and 380nm LED excitation wavelengths with emission measured at 520nm on a Zeiss Colibri LED fluorescence microscope equipped with a high speed Mrm CCD camera. Videos were recorded using the Axiovision Physiology Ca^2+^ imaging package which was also used for preparation and analysis of the ratiometric Fura 2 Ca^2+^ traces. Regions of interest (ROIs) were drawn over the whole cell and a minimum of 20 cells were analysed per experimental repeat.

### Lipid cytochemistry

Intracellular cholesterol and ganglioside GM1 localisation and accumulation were measured in cells fixed for 10 minutes with 4% paraformaldehyde (Fisher Scientific, Loughborough, UK) using filipin and fluorescein isothiocyanate (FITC) labelled cholera toxin B subunit (FITC-CtxB) respectively, as previously described^2,6^. Briefly, for cholesterol staining, fixed cells were incubated with 175μg/ml filipin in DMEM with 10% FBS at room temperature for 30mins followed by one wash in DMEM with 10% FBS and two washes in Dulbecco’s Phosphate Buffered Saline (DPBS) prior to mounting in a solution of 4.8% MOWIOL 4-88 (Merck, Darmstadt, Germany) and 12% glycerol in PBS. For ganglioside GM1 staining, fixed cells were incubated with 1μg/ml FITC-CtxB in DPBS containing 1% bovine serum albumin (BSA) and 0.1% saponin at 4**°**C for 16h followed by three 5 min washes in DPBS and mounting in MOWIOL 4-88. Lysotracker Red (Invitrogen) staining for live visualisation of lysosomes in cells was performed as described^6^, cells grown on chamberslides were incubated with 200nM Lysotracker Red in DPBS for 15min at room temperature followed by three washes in DPBS and immediate imaging. Nuclei were counterstained by addition of 2μg/ml Hoechst 33358 or Sytox Green (Invitrogen) either in PBS or added to the lysotracker staining solution. In all cases, cells were imaged using a Zeiss Colibri LED microscope with Axiovision 4.8 software, post-analysis was performed using ImageJ (NIH) and Adobe Photoshop CS6.

### Texas Red dextran endocytosis

Fluid phase endocytosis of Texas Red dextran was measured in live cells incubated for 4h with the curcumin nanoformulations in conjunction with 0.25mg/ml 10kDa Texas Red dextran in DMEM medium supplemented with 10% FBS and 1% L-glutamine. Cells were then washed 3 x 5min with cold DMEM medium with 10% FBS supplemented with 1% BSA and 0.5mg/ml unlabeled 10kDa dextran to remove non-internalised Texas Red dextran not internalized into the cells that is associated with the plasma membrane. Cells were then washed three times with DPBS and were immediately imaged using a Zeiss Colibri LED microscope.

### Thin layer chromatography (TLC)

Lipids were extracted from 100mg of the powdered curcumin nanoformulations in chloroform/methanol (VWR, West Sussex, UK) at 1:2 (v:v) as previously described^3^. After centrifugation (5 min, 1000 rpm) to remove sediment, solvent extracted lipids were washed three times by phase separation via the addition of 1mL of CHCl_3_ and 1mL of PBS, centrifugation and repeated removal of the aqueous phase, the final organic phase was dried under N_2_. Purified lipids were resuspended in ethanol, separated on high performance thin layer silica gel 60 chromatography (HPTLC) plates (Merck) and visualised with p-anisaldehyde as previously described^24^. The developing solvent systems used were: chloroform:methanol:H_2_O 65:25:4 for improved separation of phospholipids and 80:10:1 for improved separation of cholesterol from ceramides. Lipid standards were included on all TLC plates at the indicated concentrations and were obtained from Avanti Polar Lipids (Alabama, USA).

### Cell viability assays

Cellular viability following either 16h or 48h treatment with the curcumin formulations was determined on live cells by fluorescence microscopy using the early apoptotic marker Annexin A5. Following curcumin treatment, cells were incubated for 30mins on ice with 5μg/ml Alexa Fluor 488-Annexin A5 (Invitrogen) in HBSS supplemented with 10mM HEPES (pH 7.2), 1mM MgCl_2_ and 1.2mM CaCl_2_. Cells were then washed three times and imaged in the same buffer at 4°C (to prevent internalization of the Annexin A5) or by MTS assay (Promega) in 96 well plates respectively.

### Nanoparticle size analysis

The size distribution and concentration of nanoparticles in the curcumin formulations were analysed using a NanoSight. Following solubilisation in DMSO, the curcumin formulations were diluted 1 in 50,000 in mqH_2_O prior to loading onto the NanoSight and pump assisted flow over a 488 laser at a speed of 50. Movies were captured at 25FPS using a sCMOS camera and analysis was performed using the NanoSight software at a detection threshold of 2.

## Results

For this study we compared a range of curcumin formulations all of which, apart from turmeric root extract (TRE), are complexed with a range of lipid carriers in order to improve bioavailability (Table 1). Almost all, apart from BCM95s, which is solubilised in the essential oils of turmeric, contain one or a mixture of the following; lecithin, fatty acids, triglycerides, phospholipids and stearic acid. One, CGM, contains fenugreek galactomannans. We have performed the first comparison of the size properties of these curcumin nanoformulations and have confirmed that all are nanoparticles (Fig. 1a). Four, including TRE, CGM, SLN_M_ and SLN_A_ are in the range of 75-85nm whilst three have peak sizes at 100-110nm including SLN_L_, BCM95_s_ and BCM95_g_. A second broader peak ranging from 110-175nm also exists for SLN_L_, SLN_M_ and BCM95_s_ representing aggregation of these particles. A third peak at 220nm can also be seen for BCM95_s_ indicating this formulation made from the essential oils of turmeric has the most diversity in terms of particle size which is tempered by the addition of lecithin and beeswax in the case of BCM95_g_ (Table 1).

In addition to confirming that these curcumin formulations are nanoparticles we also determined the curcuminoid content in the nanoformulations via solvent extraction and separation by HPTLC as described in materials and methods. Using this method we were able to separate the three major curcuminoids, namely curcumin (largest band), desmethoxycurcumin and bis-desmethoxycurcumin in all of the nanoformulations (Fig. 1b). Our results are largely in keeping with the total curcumin content as reported (Table 1) with TRE having the highest overall curcuminoid content followed by BCM95_g_ and BCM95_s_ whereas SLN_L_, SLN_A_ and SLN_M_ have the lowest curcumin content.

In order to determine whether the curcumin supplements were likely to have any beneficial effect on NPC cells we first determined whether they could induce an elevation in cytosolic Ca^2+^ levels. All of the supplements were able to induce rapid elevation in cytosolic Ca^2+^ in both *Npc1^+/+^* and *Npc1^-/-^* astrocytes at 30μM (Fig. 2a). Interestingly, whilst the TRE, BCM95_s_, and BCM95_g_ supplements all released similar levels of Ca^2+^ in *Npc1^+/+^* and *Npc1^-/-^* astrocytes, the CGM, SLN_L_, SLN_M_, and SLN_A_ supplements all induced greater release in the NPC1 disease cells compared to controls (Fig. 2b). The SLN_A_ formulation elevated intracellular Ca^2+^ by 2-2.5 times more in the *Npc1^-/-^* cells compared to the *Npc1^+/+^* controls. Presumably this is a combined result of the ability of the SLN lipid vehicle in particular to enhance delivery across the plasma membrane coupled to altered NPC1 disease membrane fluidity^25^ leading to greater release of curcumin within the cell.

**Figure 2.**
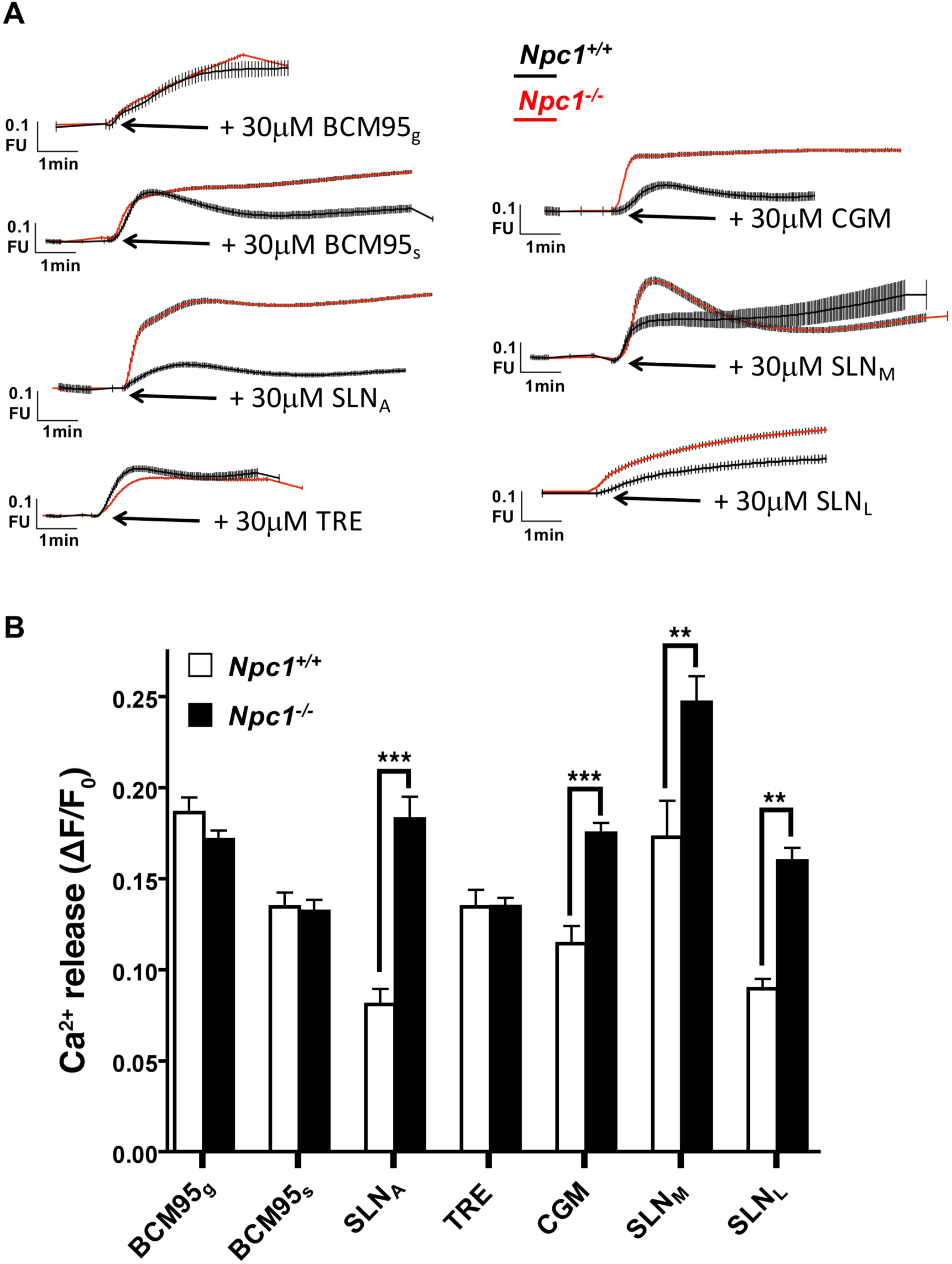
Curcumin nanoformulations all elevate cytosolic Ca^2+^ in wild-type and NPC1 disease astrocytes. (A) Averaged ratiometric traces from a single representative experiment (>30 cells analysed) illustrating elevation in cytosolic Ca^2+^, measured with Fura 2 (ΔF/F_0_), in response to 30μM addition of the indicated curcumin supplement. (B) Quantification of changes in cytosolic Ca^2+^ induced by the curcumin supplements, from 4-5 independent experiments (>100 cells). Data are given as averages ± SEM. **, p<0.01, ***, P<0.001.

Having confirmed that all the curcumin formulations were capable of elevating intracellular Ca^2+^ levels we next determined whether this could induce a reduction in NPC1 lysosomal lipid storage as previously reported with pure unformulated curcumin^2,10^. Surprisingly, although SLN_M_ had the greatest effect on elevating cytosolic Ca^2+^ in *Npc1^-/-^* disease astrocytes it had no beneficial effect on lysosomal storage. Indeed, we observed an increase in lysosomal accumulation of cholesterol (Fig. 3a), a smaller increase in ganglioside GM1 (Fig. 3b), and further expansion of the lysosomal system visualized by lysotracker staining (Fig. 3c) in *Npc1^-/-^* cells treated with SLN_M_. Despite their ability to elevate cytosolic Ca^2+^ to a greater degree in the *Npc1^-/-^* astrocytes, CGM had no effect on lipid storage or lysosomal expansion whereas SLN_L_, in a manner similar to SLN_M_, consistently led to an increase in lipid storage of cholesterol (Fig. 3a), gangliosides (Fig. 3b), and an expansion of lysosomes (Fig. 3c), no changes were observed in the *Npc1^+/+^* cells (not shown). In contrast to the other two SLN nanoformulations, SLN_A_ had no effect on cholesterol storage (Fig. 3a), ganglioside storage (Fig. 3b) or lysosomal expansion (Fig. 3c). Two curcumin supplements consistently emerged as having the greatest impact on lowering lysosomal lipid storage in the *Npc1^-/-^* cells, namely BCM95s and TRE with reduction in cholesterol (Fig. 3a), ganglioside GM1 (Fig. 3b), and lysosomal expansion (Fig. 3c) observed with BCM95s and a reduction in cholesterol (Fig. 3a) and lysosomal expansion (Fig. 3c) with TRE. No detrimental effect of any of the supplements on inducing lysosomal storage of these lipids in *Npc1^+/+^* cells was observed (not shown).

**Figure 3.**
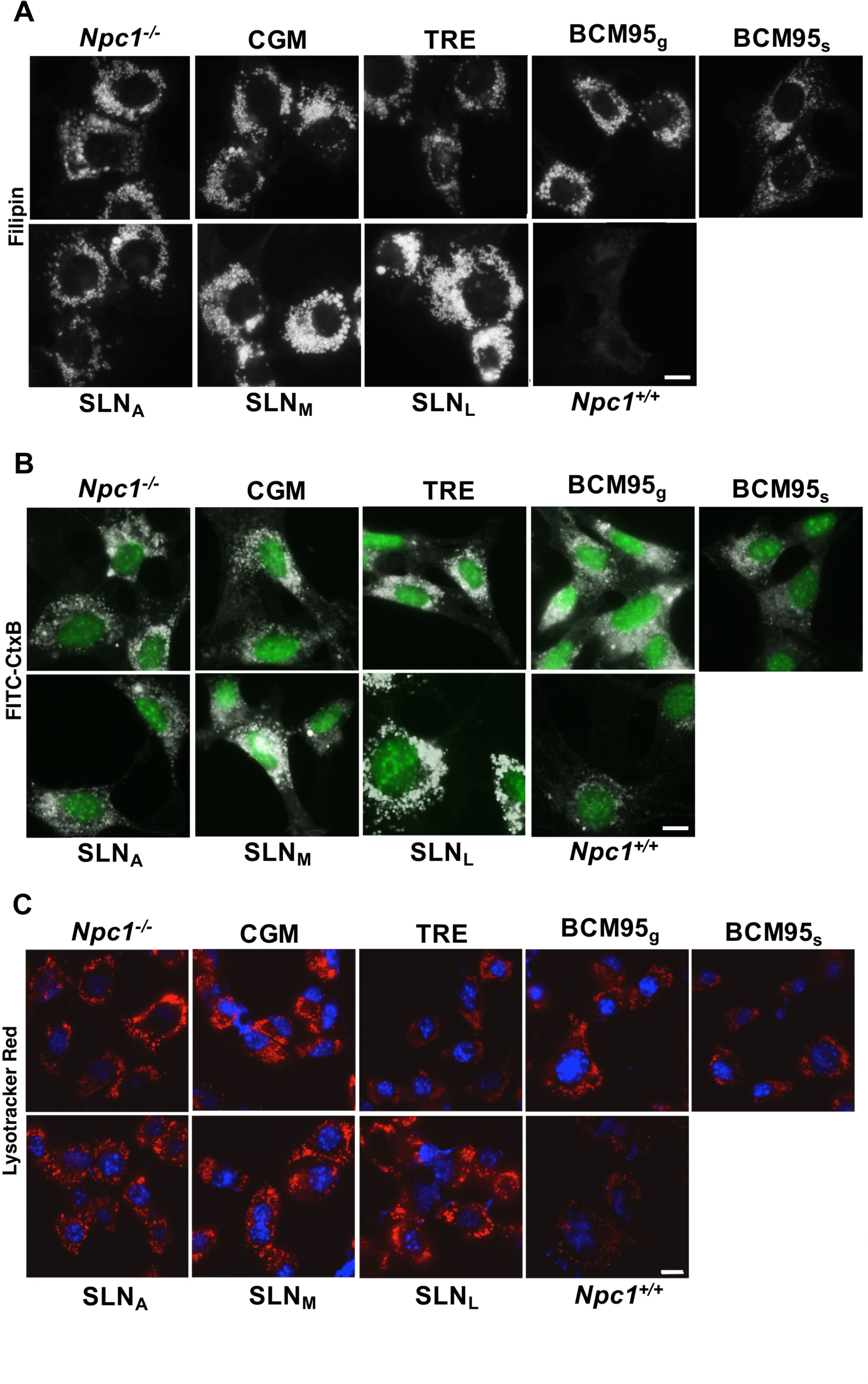
Varying effects of curcumin nanoformulations on NPC1 disease lysosomal lipid storage. Representative images from 5 independent repeats show the effects of 30μM curcumin nanoformulations on *Npc1^-/-^* astrocyte lipid storage phenotypes following a 4h (filipin, lysotracker) and 12h (FITC-CtxB) incubation (untreated *Npc1^+/+^* are included for comparison). (A) cholesterol, visualised with filipin, (B) ganglioside GM1, visualised with FITC-CtxB, (C) lysosomal expansion visualised with Lysotracker Red, (nuclei counterstained with sytox green or Hoechst 33342). Scale bar = 10μm.

To determine the cause of the elevated lipid storage levels in the *Npc1^-/-^* cells treated with SLN_L_ and SLN_M_ we investigated whether incubation with these curcumin nanoformulations had any effect on endocytosis, which is known to be altered in NPC disease and is the main route for bulk lipid entry into the cell. Following a joint incubation of the cells with both the curcumin nanoformulation and 10kDa texas red dextran for 4h we observed some key differences between the different formulations. In parallel with the reduced lipid storage observed in *Npc1^-/-^* cells treated with TRE and BCM95_s_ (Fig. 3), and in keeping with previous data on curcumin^2^ we also observed a partial correction in the endocytic transport defect (Fig. 4) with these two curcumin formulations. NPC disease cells have been shown to have a delay in transport between early and late endosomes^2,6,26^, following 4h treatment with Texas Red dextran, this probe can be seen to cluster around the nucleus in late endosomes and lysosomes in the *Npc1^+/+^* cells (Fig. 4), whereas in the *Npc1^-/-^* cells it has a broader distribution representative of early endosomes as well as some late endosomes. Both TRE and BCM95_s_, as well as SLN_A_, appear to have partially rescued this transport defect with Texas Red dextran staining now clustered in a peri-nuclear region indicative of late endosomes and lysosomes, very little staining in proximity to the plasma membrane, indicative of early endosomes, can be seen. Interestingly, both SLN_M_ and SLN_L_ appear to have either reduced the entry of texas red dextran into the *Npc1^-/-^* astrocytes or enhanced it’s recycling out of the cell as the total level of cellular fluorescence is lower by ∼65% and ∼85% respectively compared to the untreated cells (Fig. 4). This would appear to suggest a connection between the elevated lipid storage levels and a further defect in endocytosis in the *Npc1^-/-^* cells, however, we also observed reduced fluorescence indicating reduced internalization of the Texas Red dextran probe in the *Npc1^-/-^* cells treated with BCM95g (∼88%) and the *Npc1^-/-^* cells treated with CGM (∼65%). As no lipid storage was observed in these cells (Fig. 3) it must be concluded that the defect in endocytosis of texas red dextran is not the cause of the elevated lipid storage observed with SLN_M_ and SLN_L_.

**Figure 4.**
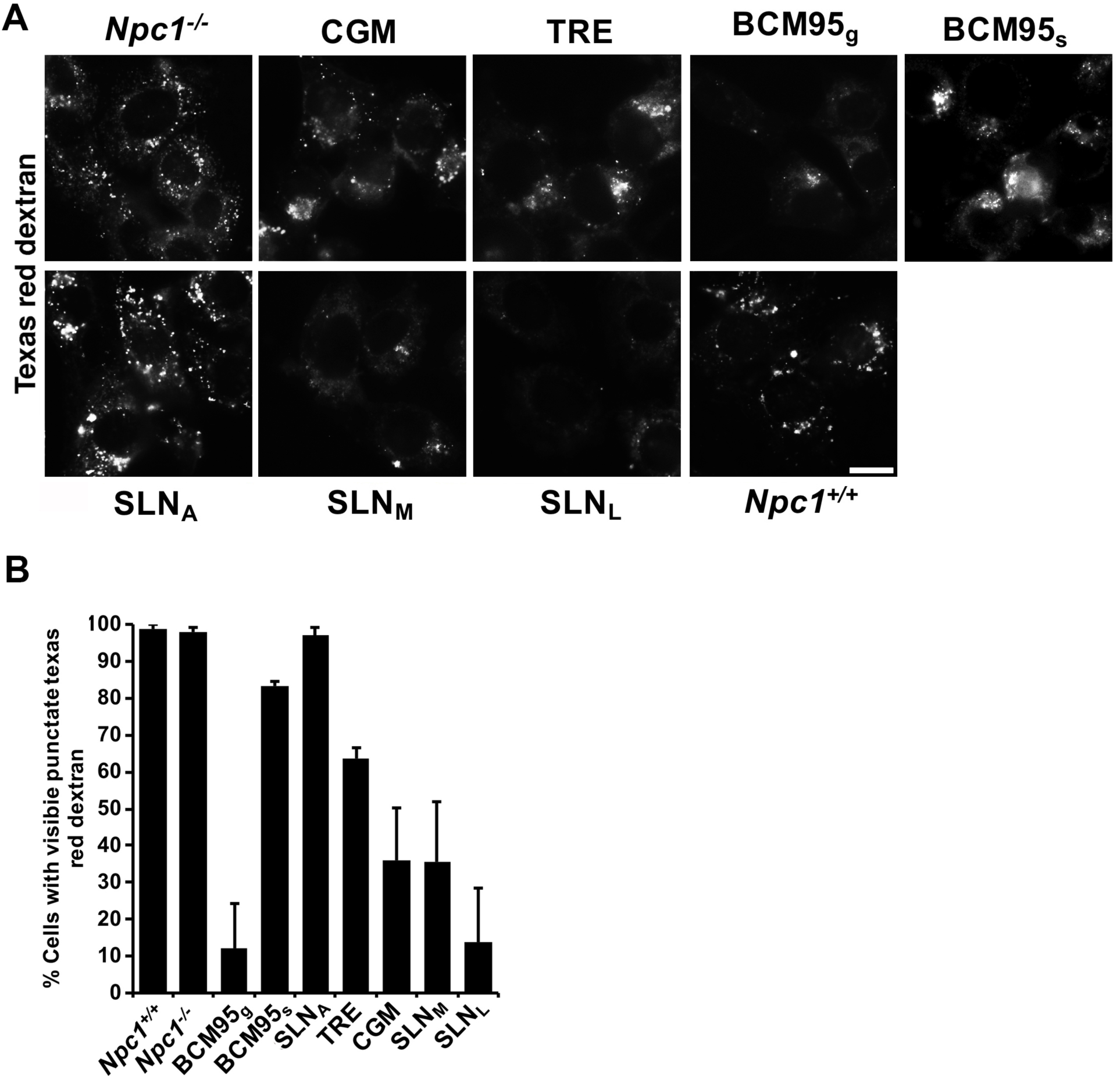
Variable effect of curcumin nanoformulations on endocytosis. (A) Representative images from 3-4 independent repeats show the effects of 30μM curcumin supplements on endocytosis of 0.25mg/ml Texas Red dextran after a 4h incubation in *Npc1^-/-^* astrocytes. (B) Analysis of the data represented in (A) whereby the number of punctate cells versus no staining was quantified across all experiments. Scale bar = 10μm.

Having observed that some of the curcumin nanoformulations increased lipid storage in NPC disease cells and that this may not be due to defects in endocytosis induced by the curcumin formulations we decided to determine the nature of the lipid species in each formulation by solvent extraction and separation by HPTLC. Perhaps unsurprisingly, TRE had the lowest lipid content with very few bands present which correlate with those observed at a higher level in BCM95_s_ and BCM95_g_ (Fig. 5a and 5b), both of which contain essential oils of curcumin that are presumably present in lower concentrations in a total turmeric root extract (TRE). CGM, which has few lipids, also contains one of these bands, the identity of which is currently unknown but could possibly be related to the galactomannan present in CGM or is a component of the curcumin used in manufacturing CGM. Interestingly, the three curcumin species are visible in all lanes (compare with Fig. 1b) between the glucosylceramide and cholesterol bands (Fig. 5a). As well as TRE and CGM, two other curcumin nanoformulations, BCM95_s_ and SLN_A_, contained very few lipids with only the reported triglycerides and fatty acids present for SLN_A_ (the band above cholesterol). Otherwise, the remaining curcumin supplements (SLN_L_, BCM95_g_, and SLN_M_) had significant levels of a variety of lipids. SLN_M_ has the highest lipid content (Fig. 5a and 5b) and is reported as using phospholipid to solubilize curcumin, by similarity to the standard this could represent lecithin (Fig. 5c) as well as to BCM95_g_ (Fig. 5a, 5b and 5c) which incorporates lecithin (Table 1). SLN_L_ is also reported to contain lecithin (Table 1), this appears to be the case although there are fewer bands when compared to the standard (Fig. 5c) with one or two fainter additional bands also present which are also seen in SLN_M_ (Fig. 5c). For the purposes of this study, our TLC analysis largely confirms the stated lipid content of these formulations whilst also indicating the presence of a few other lipids. Perhaps of most importance is that none of the supplements contained lipids that are known to accumulate in NPC disease. To confirm this we ran the HPTLC plates in two different solvent systems (Fig. 5a and 5b). First we used a solvent system comprising CHCl_3_:MeOH:H_2_O 65:25:4 to separate LBPA from sphingomyelin and which also allows visualization of neutral glycosphingolipids such as glucosylceramide (three lipids that are stored in NPC disease). We observed a small amount of LBPA and sphingomyelin in SLN_M_ but did not observe any of these lipids in the other formulations (Fig. 5a). In order to separate cholesterol and ceramide we used a solvent system comprising CHCl_3_:MeOH:H_2_O 80:10:1. We did not observe any appreciable amount of cholesterol in any of the formulations (Fig. 5b). We can therefore rule out that the lipid formulations are themselves the source of the additional lipid storage that we observe in the *Npc1^-/-^* astrocytes treated with SLN_L_ and SLN_M_. However, both SLN_L_ and SLN_M_ contain high amounts of phospholipid (Fig. 5a-c), so a change in *Npc1^-/-^* lysosomal metabolism as a result of phospholipid accumulation cannot be ruled out.

**Figure 5.**
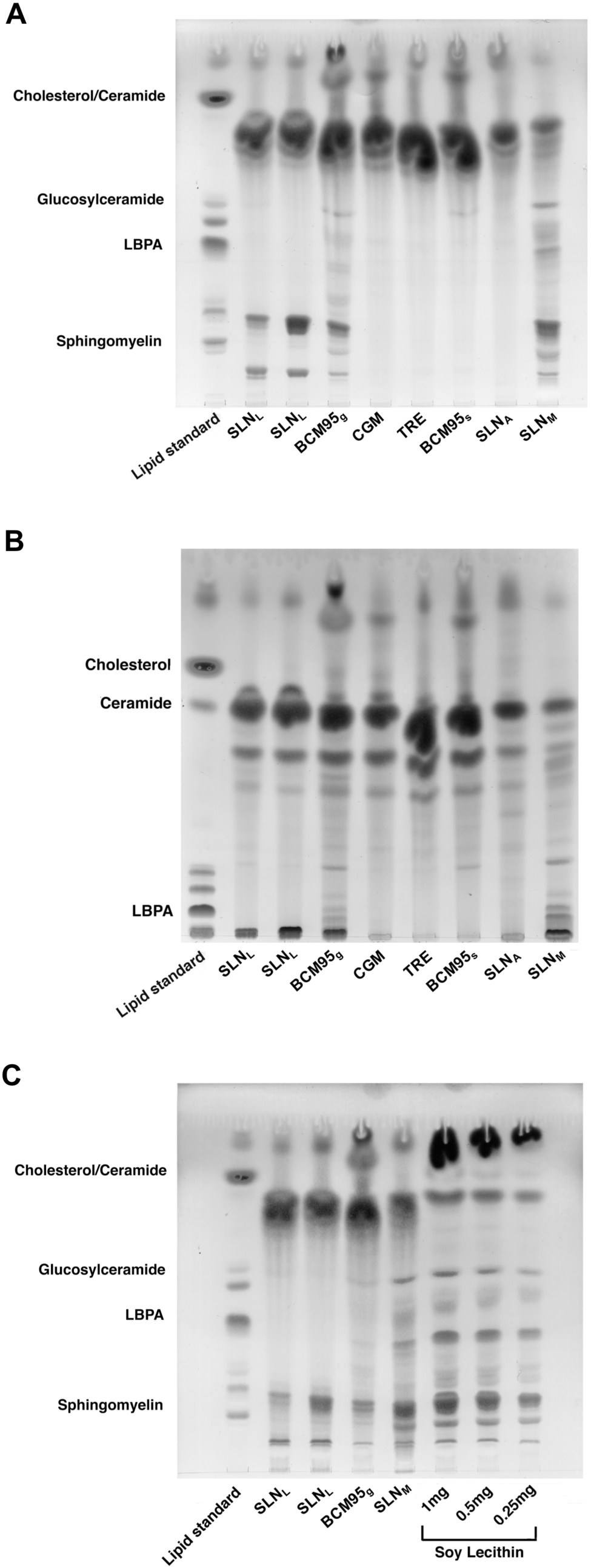
Thin layer chromatography analysis of solvent extracted curcumin nanoformulations. The lipid content of each curcumin supplement was visualised with p-anisaldehyde, A, separation of phospholipids, B, separation of ceramides and sterols. C, comparison of selected nanoformulations alongside lecithin standards. Each TLC is a representative image indicating the total lipid content extracted from 250mg of formulated curcumin, bands corresponding to lipid standards (15μg) are labelled on the y-axes, n=2.

Finally, having shown that some of the supplements could alter Ca^2+^ levels differentially between the *Npc1^+/+^* and the *Npc1^-/-^* astrocytes, with greater release in *Npc1^-/-^* (Fig. 2), that some caused an increase in lysosomal storage (Fig. 3), and that some induced defects in endocytosis (Fig. 4) potentially caused by membrane damage triggered by the nanoformulation itself^27^, we decided to test whether any of the curcumin formulations had any effect on cellular viability. First we utilized an early marker of apoptosis, extracellular live binding of FITC-Annexin A5 to plasma membrane phosphatidylserine (PS). PS is externalized to the outer leaflet of the plasma membrane as one of the first events in apoptosis. Following 16h treatment with the supplements, no staining of extracellular PS by FITC-Annexin A5 is detected in any of the conditions apart from the *Npc1^-/-^* cells treated with CGM and SLN_L_ (Fig. 6a). As a positive control to confirm that staining is indicative of apoptosis we treated cells with nigericin, a molecular poison, and observed plasma membrane FITC-Annexin A5 staining (Fig. 6a). Interestingly, some intracellular staining of FITC-Annexin A5 indicative of endosomes is observed in the *Npc1^-/-^* disease cells treated with SLN_L_ (Fig. 6a), suggesting either the possibility of necrosis or that the curcumin nanoformulation has disrupted the plasma membrane sufficiently to allow Annexin A5 to enter but not a 10kDa dextran (Fig. 4a). To confirm our viability findings with FITC-Annexin A5 we used a metabolic marker of cellular viability, namely mitochondrial activity measured using MTS. Of the curcumin formulations tested, all bar SLN_A_ had some effect on *Npc1^-/-^* mitochondrial activity and cellular viability following a 30h incubation with the supplements (Fig. 6b). TRE, BCM95_g_ and BCM95_s_ had a minimal ∼7-8% reduction in mitochondrial function whereas SLN_M_, SLN_L_ and CGM substantially, and significantly, reduced cellular viability by ∼40-50% respectively. No detrimental effect on cell viability of any of these formulations was observed on *Npc1^+/+^* cells (not shown).

**Figure 6.**
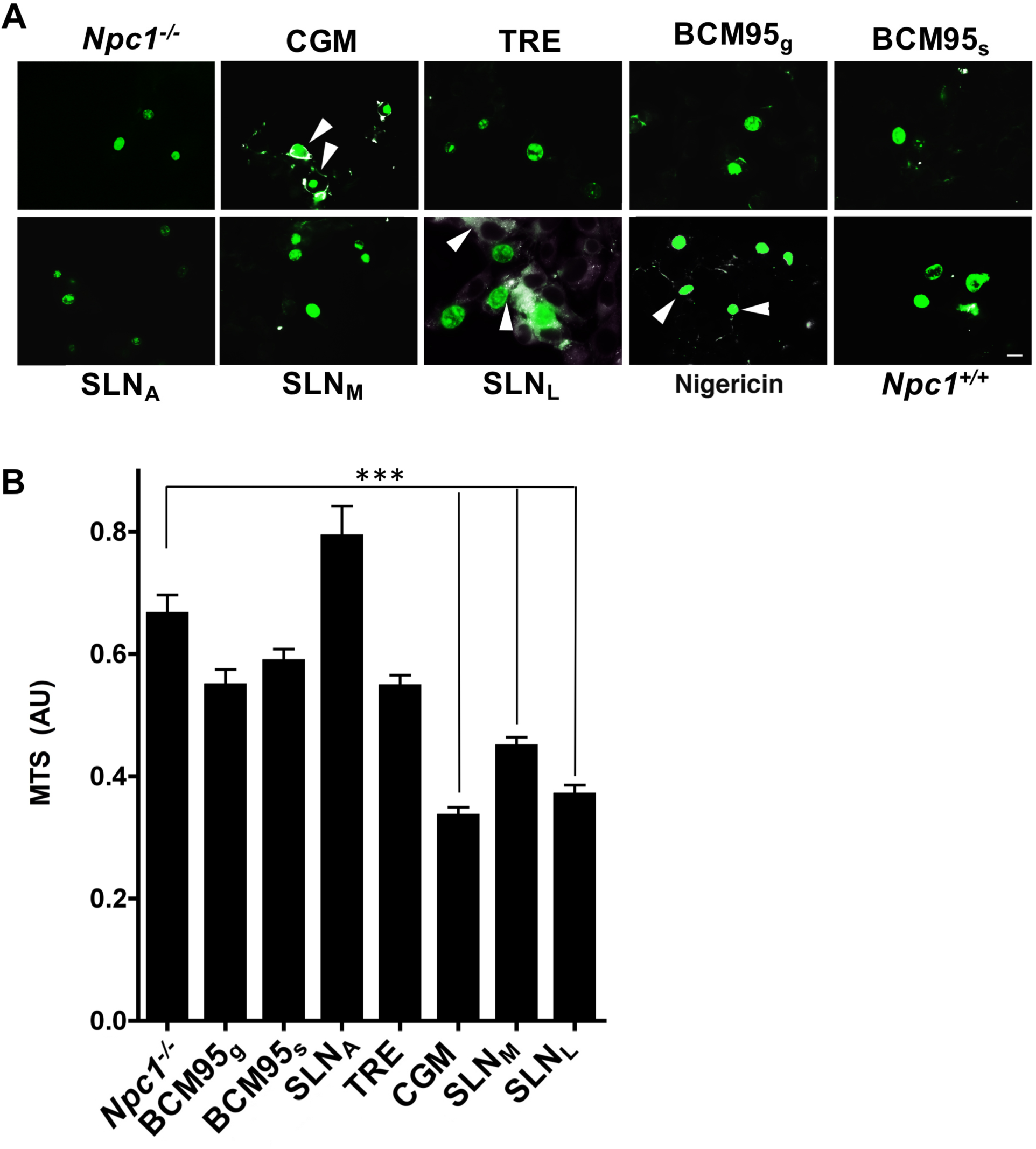
Effect of curcumin nanoformulations on NPC1 disease cellular viability. (A) Effect of 16h treatment with 30μM curcumin supplement on triggering of apoptosis in *Npc1^-/-^* astrocytes measured using Alexa Fluor 488-Annexin A5 (white) and counter stained with Hoechst 33342 (pseudocoloured in green). White arrows indicate examples of apoptotic cells with plasma membrane staining. (B) Effect of 48h treatment with 30μM curcumin supplements on mitochondrial activity in *Npc1^-/-^* astrocytes measured by MTS (absorbance units, AU). Data are given as averages ± SEM, n = 3-5. *, p<0.05, **, P<0.01, ***, P<0.001. Scale bar = 10μm.

## Discussion

Unformulated curcumin has been shown to be able to ameliorate NPC disease in five separate studies^2,13,14,28,29^, whilst curcumin formulated into a lipidated vector has been shown to have little to no benefit^13^. The aim of the present study was therefore to determine whether formulated lipidated curcumin could mediate any benefit in NPC1 disease cells. Our project focused on a range of lipidated curcumin nanoformulations (Table 1) which are used for increasing the bioavailability of curcumin which is otherwise poorly absorbed through the intestines^15^. Formulation of curcumin into a lipid vector allows for improved delivery across the blood-gut barrier, the achievement of higher concentrations of curcumin in the blood and tissues, and reduced renal clearance, all of which is essential for treating disease^19^. Recent evidence has shown that formulation into nanoparticles allows for even greater penetration into the blood and higher steady state levels, dependent on particle size^19^. Our study is the first to demonstrate that all of these formulated lipidated curcumin particles are nanoparticles, ranging in size from 50-250nm.

With respect to the ability of these nanoformulations to modulate intracellular Ca^2+^, all were capable of inducing Ca^2+^ release from the ER and elevating cytosolic Ca^2+^ levels. It was interesting to note that four, CGM, SLN_A_, SLN_M_ and SLN_L_, induced greater Ca^2+^ elevation in the *Npc1^-/-^* cells but this did not correspond with a reduction in lysosomal storage as would have been expected^2^. However, this enhanced intracellular Ca^2+^ release elicited by the majority of these nanoformulations (with the exception of SLN_A_) did correspond with increased cellular toxicity, namely reduced mitochondrial activity and apoptosis. However, another possibility is that CGM, SLN_A_, SLN_M_ and SLN_L_ may be the least stable of the nanoformulations and release their curcumin cargo in a more rapid manner upon entry into the cell compared to the other nanoformulations^15^, thus having a more significant effect on intracellular Ca^2+^.

A surprising result of our work is that although all of the nanoformulations of curcumin could modulate intracellular Ca^2+^, very few actually had an impact on NPC disease lysosomal lipid storage. As unformulated curcumin has been shown in several studies to be effective^2,14,28,29^, we hypothesised that this was due in some way to the properties of the lipid carrier. Although SLN_A_ appears to have a small benefit on some components of NPC1 lysosomal storage, no effect is seen with CGM, whilst treatment with SLN_M_ and SLN_L_ led to an elevation in lysosomal storage. This worsening of the NPC1 lysosomal storage phenotype was greatest with SLN_L_, but the exact reasons underlying this unexpected phenotype are unclear. SLN_L_ and SLN_M_ both appeared to significantly reduce endocytosis of Texas Red dextran, which might explain the elevated lysosomal storage. However, we also observed this endocytosis defect with CGM and BCM95g, neither of which had any effect on lysosomal storage, which rules this out as a possibility. Based on the similar properties of the SLN_L_ and SLN_M_ particles one possibility is that their lipid content is related to the enhanced lysosomal storage we observed in the *Npc1^-/-^* cells treated with these nanoformulations. This is supported by the lack of effect of SLN_A_, which has a similar formulation but is substantially different in that it contains sodium alginate, which can restrict the diffusion of phospholipids^30^ and may therefore ameliorate its effects on the *Npc1^-/-^* cell. However, we did not observe the presence of any NPC disease storage lipids in these nanoformulations, ruling this out as a possible cause of the elevated storage. One further potential cause of the elevated lipid storage is that the curcumin itself is trapping cholesterol within the NPC disease lysosome. Curcumin has been suggested to be capable of interacting with cholesterol^31^ and as such, delivery of curcumin into the endocytic system as a nanoparticle could potentially result in the entrapment of cholesterol within these compartments that would reduce any benefit of the elevated intracellular Ca^2+^. This mechanism could explain why only a small number of the curcumin nanoformulations led to any observable benefit. Another possibility underlying the increased lipid storage is that three of these four nanoformulations, CGM, SLN_M_ and SLN_L,_ have elicited some toxicity in the cells, as observed in Fig. 6. The associated cellular stress would lead to reduced cellular turnover and a greater degree of lipid accumulation in the non-dividing cells, supported by the reduced mitochondrial activity (Fig. 6b). However this lipid accumulation occurs, it is clearly not beneficial for lysosomal lipid levels to be elevated in cells from a lysosomal storage disease. One additional outcome of our findings is that they argue against curcumin working to rescue NPC1 lysosomal storage by exocytosis, as has been suggested^32^, as the supplement that elevates cytosolic Ca^2+^ the most (and would therefore elicit the greatest degree of exocytosis), SLN_A_, only has a minimal effect on reducing lysosomal storage levels. Of the curcumin formulations we have tested it is those least modified with lipids that overall have had the greatest beneficial effect on reducing lysosomal lipid storage in NPC1 disease cells (BCM95_s_ and TRE). These nanoformulations were able to elevate cytosolic Ca^2+^ without inducing toxicity and were able to reduce lipid storage in *Npc1^-/-^* astrocytes as previously reported with pure curcumin^2^. Our findings are in keeping with the published data from the *Npc1^-/-^* mouse model where unmodified curcumin had the greatest effect on survival and function^2^ whereas lipidated curcumin (SLN_L_) had no benefit on *Npc1^-/-^* mouse function^13^. Although this is an *in vitro* study it is important to note that the benefit of curcumin to NPC disease comes from modulation of Ca^2+^ at the ER and not as an anti-oxidant^14^. It is therefore the effect of curcumin on the individual cells of the body that needs to be considered and as such our study provides useful insight into the potential effects of curcumin formulations on NPC disease cellular function. For example, the function of the NPC1 intestine and liver is already abnormal^22^, and it is these tissues that will be primarily affected by short term treatment with these formulations. Some may transcytose, enter the blood stream and be carried to various other organs before releasing their cargo of curcumin and lipids^23^. What impact this may have on the disease course is unknown and as the only lipidated nanoformulation of curcumin to be tested in NPC disease has been SLN_L_ in the mouse model^13^ it is clear that more work is needed to determine the safety and efficacy of these nanoformulations on NPC disease.

In conclusion, as NPC1 is a lipid storage disease the use of a lipidated vehicle may not be the best approach due to the possibility that the additional lipid load could alter metabolism or endocytosis and lead to further lipid storage as we have observed. Based on our evidence^2^, from this report and from others^13,14,28,29^ it is perhaps the least modified forms of curcumin that appear to have the greatest benefit for NPC1 disease both *in vitro* and *in vivo*. Ultimately, the utilisation of curcumin itself may not be ideal for treating NPC1 disease, owing to it’s low bioavailability^15^, other more bioavailable Ca^2+^ modulators^10,11^ may yet prove to be the most effective therapeutic approach for targeting the lysosomal Ca^2+^ dysfunction in NPC disease.

## Acknowledgements

The authors would like to acknowledge the significant support and funding for this project from the Niemann-Pick Research Foundation (NPRF) as well as the support from professional services staff within the School of Biosciences. HRK and RS were supported by a summer student bursary from the NPRF. EM was supported by a PhD studentship award from the NPRF. LJH was supported by Sport Aiding Medical Research for Kids (SPARKS) and a BBSRC DTP PhD studentship alongside an MRC *in vivo* skills award. EHC was supported by a MRC DTP PhD studentship. HWE was supported by Action Medical Research and the Henry Smith charity. KW and JG were supported by a March of Dimes Basil O’Connor Starter Scholar award given to ELE who was supported by an RCUK Fellowship, a March of Dimes Basil O’Connor Starter Scholar Award and a research grant from the Royal Society.

## References

1. Lloyd-Evans, E. & Platt, F. M. Lipids on trial: the search for the offending metabolite in Niemann-Pick type C disease. Traffic 11, 419–428, doi:10.1111/j.1600- 0854.2010.01032.x (2010).

2. Lloyd-Evans, E. et al. Niemann-Pick disease type C1 is a sphingosine storage disease that causes deregulation of lysosomal calcium. Nature medicine 14, 1247– 1255, doi:10.1038/nm.1876 (2008).

3. Vruchte, D. et al. Accumulation of glycosphingolipids in Niemann-Pick C disease disrupts endosomal transport. The Journal of biological chemistry 279, 26167– 26175, doi:10.1074/jbc.M311591200 (2004).

4. Ioannou, Y. A. Guilty until proven innocent: the case of NPC1 and cholesterol. Trends in biochemical sciences 30, 498–505, doi:10.1016/j.tibs.2005.07.007 (2005).

5. Patterson, M. C., Vecchio, D., Prady, H., Abel, L. & Wraith, J. E. Miglustat for treatment of Niemann-Pick C disease: a randomised controlled study. The Lancet. Neurology 6, 765–772, doi:10.1016/S1474-4422(07)70194-1 (2007).

6. Lachmann, R. H. et al. Treatment with miglustat reverses the lipid-trafficking defect in Niemann-Pick disease type C. Neurobiology of disease 16, 654–658, doi:10.1016/j.nbd.2004.05.002 (2004).

7. Pineda, M. et al. Miglustat in patients with Niemann-Pick disease Type C (NP-C): a multicenter observational retrospective cohort study. Molecular genetics and metabolism 98, 243–249, doi:10.1016/j.ymgme.2009.07.003 (2009).

8. Choudhury, A. et al. Rab proteins mediate Golgi transport of caveola-internalized glycosphingolipids and correct lipid trafficking in Niemann-Pick C cells. J Clin Invest 109, 1541–1550, doi:10.1172/JCI15420 (2002).

9. Ruas, M. et al. Purified TPC isoforms form NAADP receptors with distinct roles for Ca(2+) signaling and endolysosomal trafficking. Current biology : CB 20, 703–709, doi:10.1016/j.cub.2010.02.049 (2010).

10. Visentin, S. et al. The stimulation of adenosine A2A receptors ameliorates the pathological phenotype of fibroblasts from Niemann-Pick type C patients. The Journal of neuroscience : the official journal of the Society for Neuroscience 33, 15388–15393, doi:10.1523/JNEUROSCI.0558-13.2013 (2013).

11. Xu, M. et al. delta-Tocopherol reduces lipid accumulation in Niemann-Pick type C1 and Wolman cholesterol storage disorders. The Journal of biological chemistry 287, 39349–39360, doi:10.1074/jbc.M112.357707 (2012).

12. Logan-Smith, M. J., Lockyer, P. J., East, J. M. & Lee, A. G. Curcumin, a molecule that inhibits the Ca2+-ATPase of sarcoplasmic reticulum but increases the rate of accumulation of Ca2+. The Journal of biological chemistry 276, 46905–46911, doi:10.1074/jbc.M108778200 (2001).

13. Borbon, I. A. et al. Lack of efficacy of curcumin on neurodegeneration in the mouse model of Niemann-Pick C1. Pharmacology, biochemistry, and behavior 101, 125– 131, doi:10.1016/j.pbb.2011.12.009 (2012).

14. Williams, I. M. et al. Improved neuroprotection using miglustat, curcumin and ibuprofen as a triple combination therapy in Niemann-Pick disease type C1 mice. Neurobiology of disease 67, 9–17, doi:10.1016/j.nbd.2014.03.001 (2014).

15. Prasad, S., Tyagi, A. K. & Aggarwal, B. B. Recent developments in delivery, bioavailability, absorption and metabolism of curcumin: the golden pigment from golden spice. Cancer research and treatment : official journal of Korean Cancer Association 46, 2–18, doi:10.4143/crt.2014.46.1.2 (2014).

16. Begum, A. N. et al. Curcumin structure-function, bioavailability, and efficacy in models of neuroinflammation and Alzheimer’s disease. The Journal of pharmacology and experimental therapeutics 326, 196–208, doi:10.1124/jpet.108.137455 (2008).

17. Miranda, M. I., Gonzalez-Cedillo, F. J. & Diaz-Munoz, M. Intracellular calcium chelation and pharmacological SERCA inhibition of Ca2+ pump in the insular cortex differentially affect taste aversive memory formation and retrieval. Neurobiology of learning and memory 96, 192–198, doi:10.1016/j.nlm.2011.04.010 (2011).

18. Chiu, S. S. et al. Differential distribution of intravenous curcumin formulations in the rat brain. Anticancer research 31, 907–911 (2011).

19. Bi, C. et al. Particle size effect of curcumin nanosuspensions on cytotoxicity, cellular internalization, in vivo pharmacokinetics and biodistribution. Nanomedicine 13, 943–953, doi:10.1016/j.nano.2016.11.004 (2016).

20. C. W. Vockley. FYI: Issues Regarding Curcumin Therapy for NPC, http://nnpdf.org/Curcumin.html (2009).

21. Boadu, E. et al. Correction of apolipoprotein A-I-mediated lipid efflux and high density lipoprotein particle formation in human Niemann-Pick type C disease fibroblasts. The Journal of biological chemistry 281, 37081–37090, doi:10.1074/jbc.M606890200 (2006).

22. Patterson, M. C. et al. Recommendations for the diagnosis and management of Niemann-Pick disease type C: an update. Molecular genetics and metabolism 106, 330–344, doi:10.1016/j.ymgme.2012.03.012 (2012).

23. Kadam, R. S., Bourne, D. W. & Kompella, U. B. Nano-advantage in enhanced drug delivery with biodegradable nanoparticles: contribution of reduced clearance. Drug metabolism and disposition: the biological fate of chemicals 40, 1380–1388, doi:10.1124/dmd.112.044925 (2012).

24. Maue, R. A. et al. A novel mouse model of Niemann-Pick type C disease carrying a D1005G-Npc1 mutation comparable to commonly observed human mutations. Human molecular genetics 21, 730–750, doi:10.1093/hmg/ddr505 (2012).

25. Koike, T. et al. Decreased membrane fluidity and unsaturated fatty acids in Niemann-Pick disease type C fibroblasts. Biochim Biophys Acta 1406, 327–335 (1998).

26. Mayran, N., Parton, R. G. & Gruenberg, J. Annexin II regulates multivesicular endosome biogenesis in the degradation pathway of animal cells. Embo J 22, 3242– 3253 (2003).

27. Panariti, A., Miserocchi, G. & Rivolta, I. The effect of nanoparticle uptake on cellular behavior: disrupting or enabling functions? Nanotechnol Sci Appl 5, 87–100, doi:10.2147/NSA.S25515 (2012).

28. Efthymiou, A. G. et al. Rescue of an in vitro neuron phenotype identified in Niemann-Pick disease, type C1 induced pluripotent stem cell-derived neurons by modulating the WNT pathway and calcium signaling. Stem Cells Transl Med 4, 230–238, doi:10.5966/sctm.2014-0127 (2015).

29. Fineran, P. et al. Pathogenic mycobacteria achieve cellular persistence by inhibiting the Niemann-Pick Type C disease cellular pathway. Wellcome Open Res 1, 18, doi:10.12688/wellcomeopenres.10036.1 (2016).

30. Mackie, A. R. et al. Sodium alginate decreases the permeability of intestinal mucus. Food Hydrocoll 52, 749–755, doi:10.1016/j.foodhyd.2015.08.004 (2016).

31. Jourghanian, P., Ghaffari, S., Ardjmand, M., Haghighat, S. & Mohammadnejad, M. Sustained release Curcumin loaded Solid Lipid Nanoparticles. Adv Pharm Bull 6, 17–21, doi:10.15171/apb.2016.004 (2016).

32. Canfran-Duque, A. et al. Curcumin promotes exosomes/microvesicles secretion that attenuates lysosomal cholesterol traffic impairment. Molecular nutrition & food research 58, 687–697, doi:10.1002/mnfr.201300350 (2014).

